# Yeast mtDNA transcription initiation in single nucleotide addition steps

**DOI:** 10.1101/2022.09.27.509796

**Authors:** Quinten Goovaerts, Jiayu Shen, Brent De Wijngaert, Urmimala Basu, Smita S. Patel, Kalyan Das

**Affiliations:** Laboratory of Virology and Chemotherapy, Rega Institute for Medical Research, KU Leuven, 3000 Leuven, Belgium; Department of Microbiology, Immunology and Transplantation, KU Leuven, 3000 Leuven, Belgium; Department of Biochemistry and Molecular Biology, Robert Wood Johnson Medical School, Rutgers University, Piscataway, NJ 08854, USA

## Abstract

Transcription initiation catalyzed by the RNA polymerase is a multistep process involving promoter binding, transcription bubble formation, abortive RNA synthesis, and transition into elongation following promoter escape. We report cryo-EM structures of yeast mitochondrial RNA polymerase initiation complexes (ICs) with transcription factor MTF1 catalyzing RNA synthesis from *de novo* initiation to 6-mer synthesis at single-nucleotide steps on fully-resolved transcription bubbles. The growing RNA:DNA hybrid is accommodated by continuous scrunching of the template strand while the non-template and MTF1 C-tail in the polymerase cleft are structurally reorganized. Each nucleotide addition accumulates stress energy, which drives abortive RNA synthesis during early transcription initiation steps and promoter release later. The non-template scrunches as loops in IC2/IC3, and unscrunching assists abortive synthesis of 2-/3-mer RNAs. Subsequently, in IC5 and IC6, the non-template strand assumes a stable structure by stacking its bases into a spiral staircase-like structure that supports processive synthesis. In IC6, the template scrunches to the maximum and places the -1 nucleotide in a pocket near the thumb domain. Subsequently, the -1 nucleotide acts as a pivot point for promoter escape ushering the IC into the elongation phase. The structural snapshots visualize the interplay between abortive and productive synthesis regulating transcription initiation.

---

Transcription initiation is a multistep process that begins with the DNA-dependent RNA polymerase (RNAP) binding and unwinding a specific region of the promoter DNA to generate a transcription bubble with an exposed transcription start site (TSS)^1^. *De novo* RNA synthesis from TSS is initiated using NTPs, and the resulting 2-mer RNA is elongated through a DNA scrunching mechanism^2-4^. During initiation, the RNAP remains bound to the upstream promoter region, and the melting of each downstream DNA base-pair brings the template nucleotide into the polymerase cleft to engage complementing NTP for RNA synthesis. Each nucleotide addition extends the RNA:DNA hybrid and the non-template (NT) strand that expand the transcription bubble creating stressed intermediates, which result in backtracking and abortive RNA synthesis^5,6,7,8^. However, after synthesizing a critical 8-to 12-mer RNA, the RNAP undergoes conformational changes to convert the initiation complex (IC) into an elongation complex (EC). This basic mechanism of transcription initiation is conserved in all domains of life. However, the molecular complexity of the transcription machinery varies, increasing from bacteriophages to humans ^9-12^. Challenges remain in visualizing how structural changes that generate scrunched intermediate states exhibiting both on-pathway RNA synthesis and off-pathway backtracking/abortive events guide the transcription initiation process.

Mitochondrial RNAPs (mtRNAPs) are essential for the transcription and replication of the mitochondrial DNA and in generating the components responsible for energy production^13,14^. The mtRNAPs are closely related to single-subunit bacteriophage T7 RNAP^15^. The yeast (*Saccharomyces cerevisiae)* mtRNAP (y-mtRNAP; RPO41) shares structural and functional similarities with human mtRNAP (h-mtRNAP; POLRMT). Like most DNA-dependent RNAPs, mtRNAPs rely on one or more transcription factors to initiate transcription. The y-mtRNAP requires one factor, MTF1, and h-mtRNAP requires two factors, TFAM and TFB2M, for promoter-specific transcription initiation^10,11,16-18^; T7 RNAP does not require any factor. Recently, we published high-resolution structures of y-mtRNAP:MTF1 with a DNA promoter in the partially-melted initiation complex (PmIC) and a transcribing IC3 states^19^. These structures resolved the complete transcription bubble, which could not be traced in the IC structures of h-mtRNAP^20^ or T7 RNAP^21^. The PmIC structure showed that MTF1 stabilizes the initiation bubble by interacting base-specifically with a conserved promoter sequence (−4) AAG (−2) in the NT strand. The IC3 structure showed the +1 to +3 template nucleotides are base-paired with a 2-mer RNA and an incoming NTP at the polymerase active site. The NT strand was scrunched into a loop in IC3. These structures provided the basis for a systematic study to visualize stepwise DNA scrunching and structural adaptations balancing the odds of abortive versus productive RNA synthesis at each nucleotide addition step.

Here, we report single particle cryo-EM structures of four transcribing complexes of y-mtRNAP:MTF1 – IC2, IC4, IC5, and IC6, each with a fully resolved transcription bubble and an RNA:DNA hybrid. The structures from PmIC to IC6 allow us to visualize the entire transcription initiation process up to 6-mer RNA synthesis at single-nucleotide addition steps (Movies 1 and 2). The structures reveal how the proteins and the transcription bubble respond and adapt to growing RNA:DNA, and stabilize each intermediate initiation state. Using these IC structures, we could address questions such as: (i) How does DNA scrunching help accommodate the growing RNA:DNA duplex and the transcription bubble in the polymerase cleft? (ii) Do both template and NT DNA strands scrunch during transcription initiation steps, and how do these structural changes affect the initiation steps? (iii) What drives abortive synthesis over RNA elongation, and why are only short RNAs of specific length aborted? (iv) How might the conformational changes trigger the transition into elongation?

## Results

### Cryo-EM structures of IC2, IC4, IC5, and IC6

We used two pre-melted (−4 to +2) DNA scaffolds of a consensus promoter sequence with slightly different downstream sequences to generate the IC2 -IC6 states for this structural study (Fig. 1a). Y-mtRNAP and MTF1 were assembled on the two DNA promoters and the resulting PmICs were purified over a size-exclusion column^22^. Defined lengths of RNA:DNA hybrids were created *in situ* by reacting the PmIC with appropriate NTPs in the presence of Mg^2^+ ions (Fig. 1a). The IC2 was generated by incubating the PmIC with GTP, and IC4 to IC6 states were obtained by adding appropriate NTPs to a starting 2-mer RNA, pppGpG. Single-particle cryo-EM structures of IC2, IC4, IC5, and IC6 were determined at 3.47, 3.44, 3.39, and 3.62 Å resolution, respectively (Supplementary Table 1). The experimental density maps resolved all the critical structural elements, including the complete transcription bubbles (Supplementary Fig. 1) and interacting protein residues. The overall structure of y-mtRNAP:MTF1 complex on the DNA remains unaltered in all IC states (Fig. 1b, c). Together with the published PmIC (PDB: 6YMV) and IC3 (PDB: 6YMW) structures^19^, the four new IC structures help visualize the progression of transcription initiation (Fig. 1d, e, f). The subsequent transition from IC6 to EC was modeled using the h-mtRNAP EC crystal structure (PDB:4BOC)^23^. The structural states morphed into movies that show the dynamic process of transcription initiation and transition to elongation (Movies 1 and 2).

**Fig. 1.**
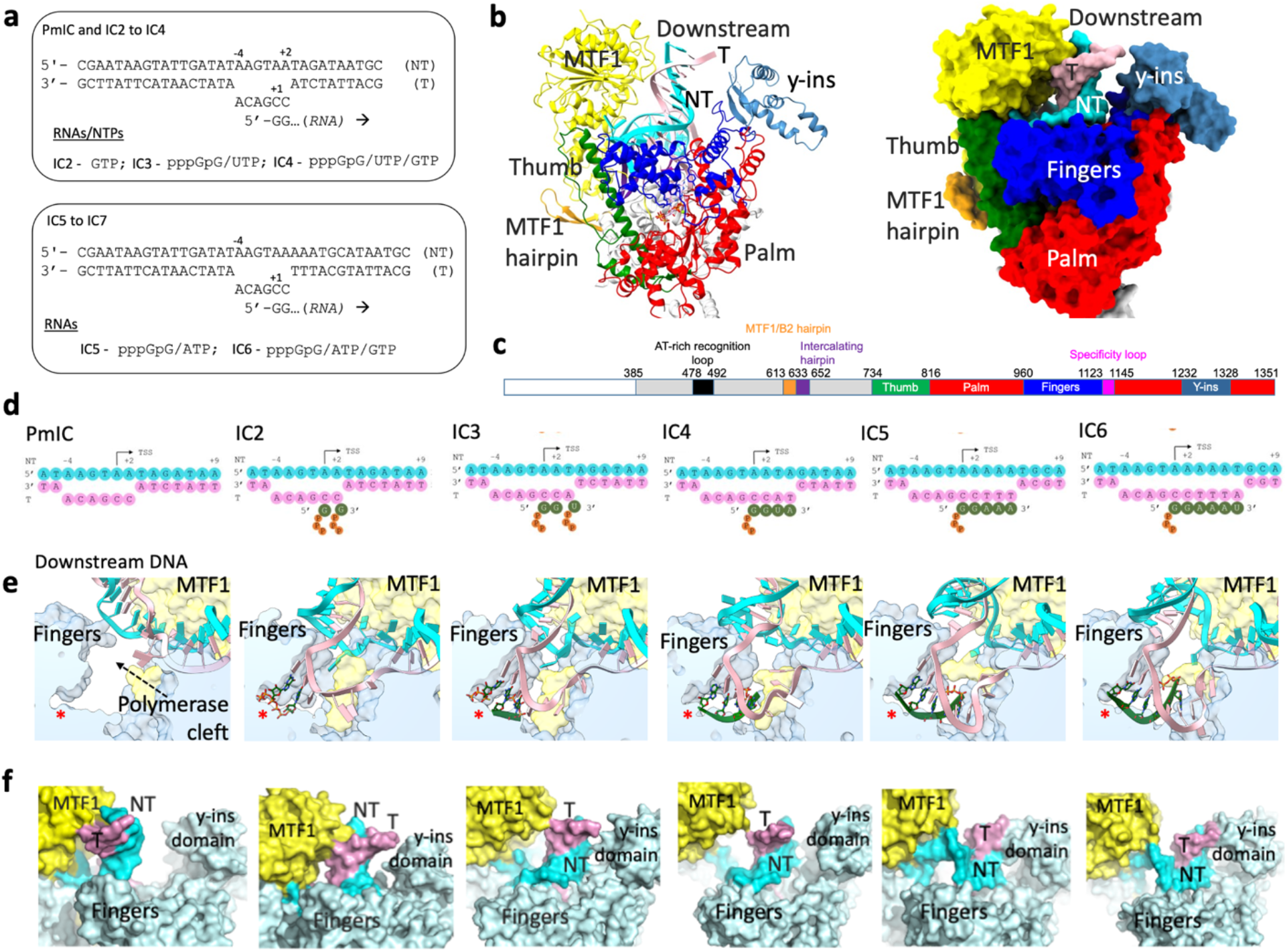
Overview of the transcription initiation states of y-mtRNAP from PmIC to IC6 at single-nucleotide addition steps. **a.** The bubble DNA promoter oligonucleotides and the NTPs used for a stepwise synthesis of RNA from 2-to 6-mer for trapping the IC2 to IC6 states with y-mtRNAP and MTF1. **b.** A cartoon (left) and surface (right) representations of the IC2 structure showing MTF1 in yellow and color-coded y-mtRNAP subdomains (fingers, blue; palm, red; thumb, green), template, pink; non-template, cyan; y-ins domain, steel blue. **c.** The color-coded 1-D representation of the structural elements of y-mtRNAP. **d.** The schematic representations of the transcription bubbles in six different states from PmIC to IC6; cyan non-template and pink template DNA, and green RNA. **e.** Zoomed views of the polymerase clefts and the transcription bubble in the PmIC and IC states; the molecular surfaces of y-mtRNAP and MTF1 are in light blue and yellow, respectively, and the active site is marked (*). **f.** Zoomed views showing the relative positioning of the downstream DNA in the “downstream cleft” defined by the space between MTF1 and y-ins; y-ins is a characteristic domain in yeast that has no T7 RNAP or h-mtRNAP homolog.

The y-mtRNAP has a right-hand-shaped C-terminal polymerase domain (CTD) with palm, fingers, and thumb subdomains present in all single-subunit RNAPs (Fig. 1b, c)^24^. These subdomains enclose the “polymerase cleft”, which contains the active site for RNA synthesis and accommodates the transcription bubble, including the RNA:DNA duplex (Fig. 1d, e). The MTF1 is positioned on the top of the polymerase cleft like a lid (Fig. 1b). The two proteins are held together primarily through two sets of interactions – (i) between the MTF1 N-terminal domain (residues 1 – 252) and the y-mtRNAP thumb tip (residues 772 – 779), and (ii) between the MTF1 C-terminal domain (aa 255 – 341) and a y-mtRNAP hairpin (aa 613 – 633), termed as “MTF1 hairpin”. The MTF1 hairpin has a homolog “B2 hairpin” in h-mtRNAP to support TFB2M binding^19,20^. The NT strand in the bubble region interacts primarily with the MTF1 and the template strand with y-mtRNAP. The consensus promoter sequence upstream of the bubble interacts with the specificity loop, AT-rich recognizing loop, and the intercalating hairpin of y-mtRNAP. The conserved (−4) AAG (−2) NT nucleotides in the bubble interact with MTF1. These interactions of the upstream promoter duplex and bubble with protein residues remain unaltered throughout initiation steps, starting from PmIC to IC6.

The downstream DNA duplex is positioned in a flexible cleft formed by the N-terminal domain of MTF1 and the y-ins (yeast RNAP insertion) domain of y-mtRNAP (Fig. 1b, f) that we refer to as the “downstream cleft”. The y-ins domain is a stretch of ∼100 amino acid residues (1232 – 1328) unique to y-mtRNAP and absent in T7 RNAP and h-mtRNAP. The positional adaptability of y-ins allows the downstream cleft to accommodate the DNA duplex in response to transcription bubble expansion (Fig. 1f). Due to the dynamic behavior of the downstream DNA and its surroundings, y-ins has a lower resolution than the core. The AlphaFold^25^ model for y-ins fitted reliably to the density with minor adjustments (Supplementary Fig. 2). The y-ins domain has a four-helix bundle and a small two-stranded anti-parallel β-sheet, and y-ins interacts nonspecifically with the downstream DNA backbone.

### IC2 and IC3 capture the NTP-bound states prior to catalysis

The IC2 intermediate captures two GTP molecules bound at the TSS and poised for catalysis. The +1 GTP is bound at the priming site (P site, also referred to as post-insertion site) and the +2 GTP at the NTP-binding site (N site, also referred to as insertion site; Fig. 2a). In the absence of +3 NTP in our experiment, we expected to trap one or more of the following states: (i) a synthesized 2-mer pppGpG RNA base-paired with the template +1 and +2 nucleotides, (ii) two GTPs bound at the +1 and +2 positions prior to catalysis, and/or (iii) the PmIC resulted after the release of the 2-mer RNA as an abortive product. However, we trapped only the GTP-bound state. The *in vitro* transcription assay shows a significant amount of GTP is converted to 2-mer RNA (Fig. 2b); i.e., the GTP-bound IC2 state is captured in the cryo-EM sample under steady-state RNA synthesis conditions. Thus, in the abortive cycle of 2-mer synthesis, the GTP-bound IC2 state must be more stable than a 2-mer RNA-bound state, which dissociates readily if the next nucleotide is unavailable. Additionally, we did not observe the coexistence of the PmIC state in the IC2 cryo-EM sample, suggesting the initiating nucleotides bind rapidly and deplete the PmIC (Fig. 2c). In contrast, the IC3 sample in our previous study captured IC3 and PmIC coexisting in solution^19^, because both 2-mer RNA + UTPαS binding and the catalysis are slow (Fig. 2C).

**Fig. 2.**
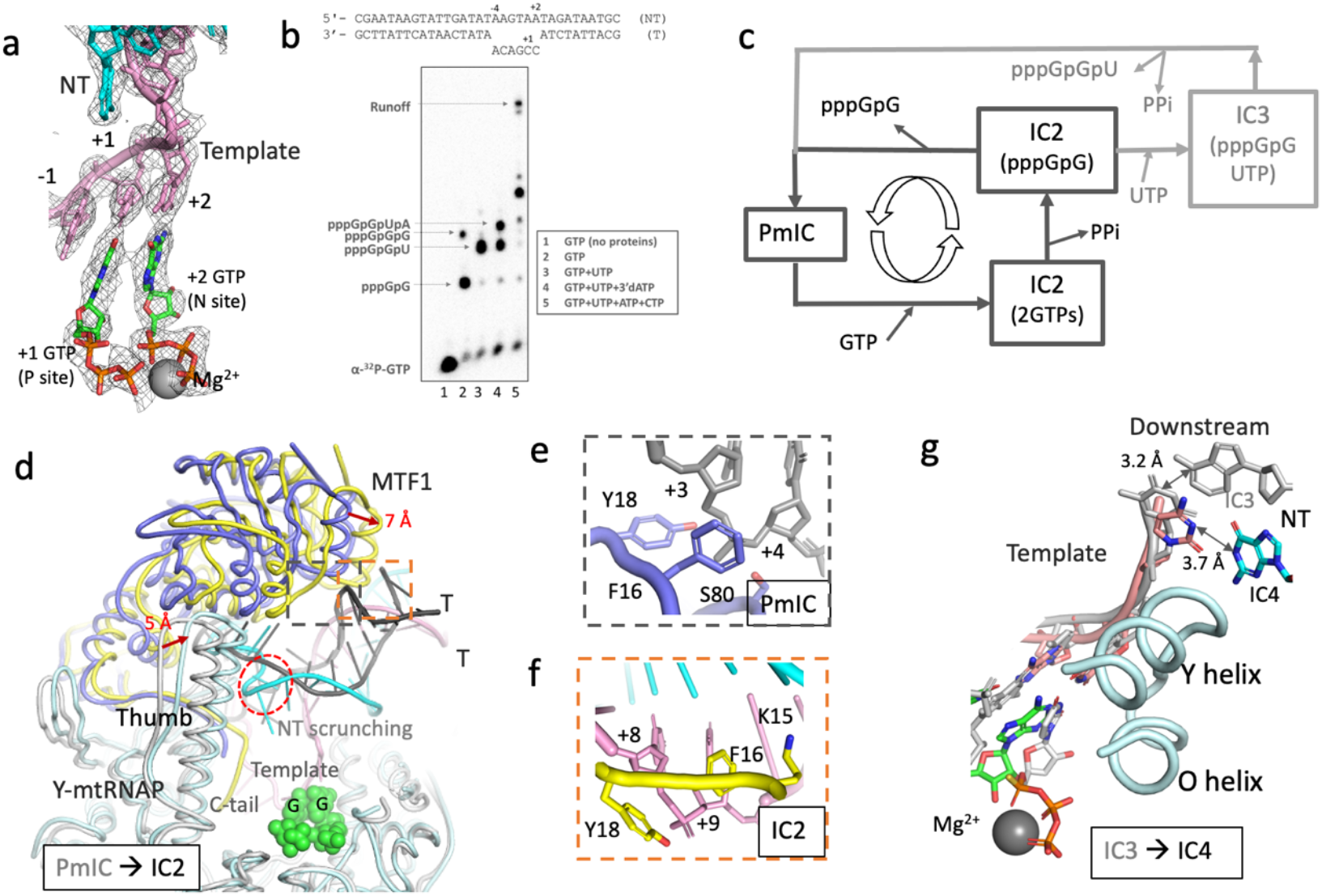
Structures of IC2 to IC4 states reveal the basis for abortive synthesis. **a**. The cryo-EM density map displayed at 2.5σ level defines the two GTP molecules at the polymerase active site poised for catalysis and base-paired with the template +1 and +2 nucleotides in IC2; NT – non-template. The template -1 base stacks with the +1 base pair. **b**. The gel image shows the RNA products from our *in vitro* transcription reactions of y-mtRNAP and MTF1 on the bubble promoter under the given NTP conditions (box). **c**. A schematic representation of IC2 and IC3 abortive cycles. The PmIC binds the initiating GTP molecules to form IC2 (2GTPs). The phosphodiester bond formation and release of pyrophosphate (PPi) convert the IC2 (2GTPs) to IC2 (pppGpG). In the absence of the next NTP, the 2-mer RNA is readily released as an abortive product, and the IC2 state returns to PmIC, which recruits new GTP molecules to continue the cyclic process. This leads to the accumulation of 2-mer pppGpG RNA as evident from the gel image in panel b. A similar abortive process occurs during IC3 formation from pppGpG and UTP. The abortive cycle among PmIC and IC3 synthesizes 3-mer RNA, pppGpGpU. The 2GTP-bound structure is trapped in IC2 sample because, the catalytic step is the slowest. The IC3 sample yield both PmIC and IC3 (pppGpG) structures because the catalytic step and loading of 2-mer RNA + UTPαS are slower compared to a fast release of 3-mer RNAs. **d**. Superposition of the IC2 (yellow MTF1, light-blue y-mtRNAP, cyan NT, and pink template) and PmIC (blue MTF1 and gray y-mtRNAP/DNA) structures shows the correlated motions of MTF1 and thumb subdomain (red arrows) to form a closed IC2 upon binding of the initiating GTP molecules (green). The scrunched NT strand is circled red. **e**. Interactions of the template strand +3 and +4 nucleotide backbone with F16/Y18 groove of MTF1 in PmIC. **f**. Interactions of the template strand +8 and +9 backbone with F16/Y18 groove of MTF1 in IC2. **g**. Cα-superposition of y-mtRNAP IC3 structure (gray) on IC4 (green RNA, pink template, cyan NT, and light blue y-mtRNAP fingers helices) shows that the 3’-end nucleotide of the RNA in IC4 has moved halfway (∼3.5 Å) towards the P site from the N site. The +5 base-pair in the downstream DNA is partially melted with the average base-pair H-bond distance of ∼3.7 Å compared to 3.2 Å for the +4 base pair in IC3 (gray).

The transition from PmIC to IC2 occurs with a large conformational change that moves the template strand by ∼22 Å, allowing the TSS (+1 and +2 nucleotides) to base-pair with two initiating GTPs at the active site (Fig. 2a, d). This conformational change of the template was observed earlier when structures of PmIC and IC3 were compared^19^. Solving the IC2 structure completes the pathway from PmIC ⟶ IC2 ⟶ IC3. Template alignment at the active site bends the downstream DNA from 60° in PmIC to 120° in IC2. To accommodate this DNA bending, the MTF1 N-terminal domain moves by ∼7 Å towards the y-ins domain, closing the cleft around the downstream DNA (Fig. 1f, 2d). Harmoniously, the tip of the y-mtRNAP thumb that interacts with MTF1 (aa 105, 153-158) moves by ∼5 Å to maintain protein-protein interactions. The combined changes from DNA bending, transcription bubble expansion, and downstream cleft closing alter the interactions of the downstream DNA with MTF1. In PmIC, the +4/+5 template backbone interacts with the F16/Y18 groove of MTF1 for stabilizing the DNA in the early events of promoter bending and melting (Fig. 2e). In IC2, the F16/Y18 groove interactions shift to the +8/+9 template backbone (Fig. 2f). The downstream cleft starts opening in IC3 and continues to open further in IC4 (Supplementary Fig. 3), with loss of F16/Y18 and downstream DNA interactions (Fig. 1f).

Unlike the NTP-bound IC2 and IC3 states, the IC4 and subsequent states are captured with fully synthesized RNAs, base-paired to the DNA template. The synthesized 4-mer RNA, pppGpGpUpA, in IC4 is base-paired to the template nucleotides +1 to +4 (Supplementary Fig. 1). No stable PmIC or other intermediate state was observed coexisting in the IC4 cryo-EM sample. Interestingly, the 4-bp duplex RNA:DNA is in a half-translocated state; i.e., the 3’-end of the nascent RNA has moved by ∼3.5 Å at the C1’ position from the N site but has not reached the P site; the distance between the C1’ positions of the nucleotides at N and P sites in IC2/IC3 is ∼6Å. Additionally, the +5 base-pair in the downstream DNA is partially melted in IC4; the average base-pair H-bond distance for the +5 G:C is ∼3.7 Å (Fig. 2g), longer than the expected ∼3 Å for a typical base-pair. Thereby, the structure of IC4 favors nucleotide incorporation over abortive synthesis.

### Non-template DNA scrunching contributes to abortive synthesis

Abortive synthesis is an off-pathway event observed during transcription initiation by all cellular RNAPs including bacterial and bacteriophage T7 RNAPs^3,26,27^. Unlike T7 and bacterial systems where the transcription is aborted at all IC states, the *in vitro* transcription assays show that y-mtRNAP:MTF1 synthesizes significant amounts of 2 -and 3-mer abortive RNAs, after which RNA synthesis becomes relatively processive (Fig. 3a). Quantification of the gel bands finds that ∼46% and ∼36% products are 2 -and 3-mer, respectively, in comparison to ∼14% runoff product; the 4 -and 5-mer are only ∼2% and <1%, respectively, even though all NTPs were present in the assay.

**Fig. 3.**
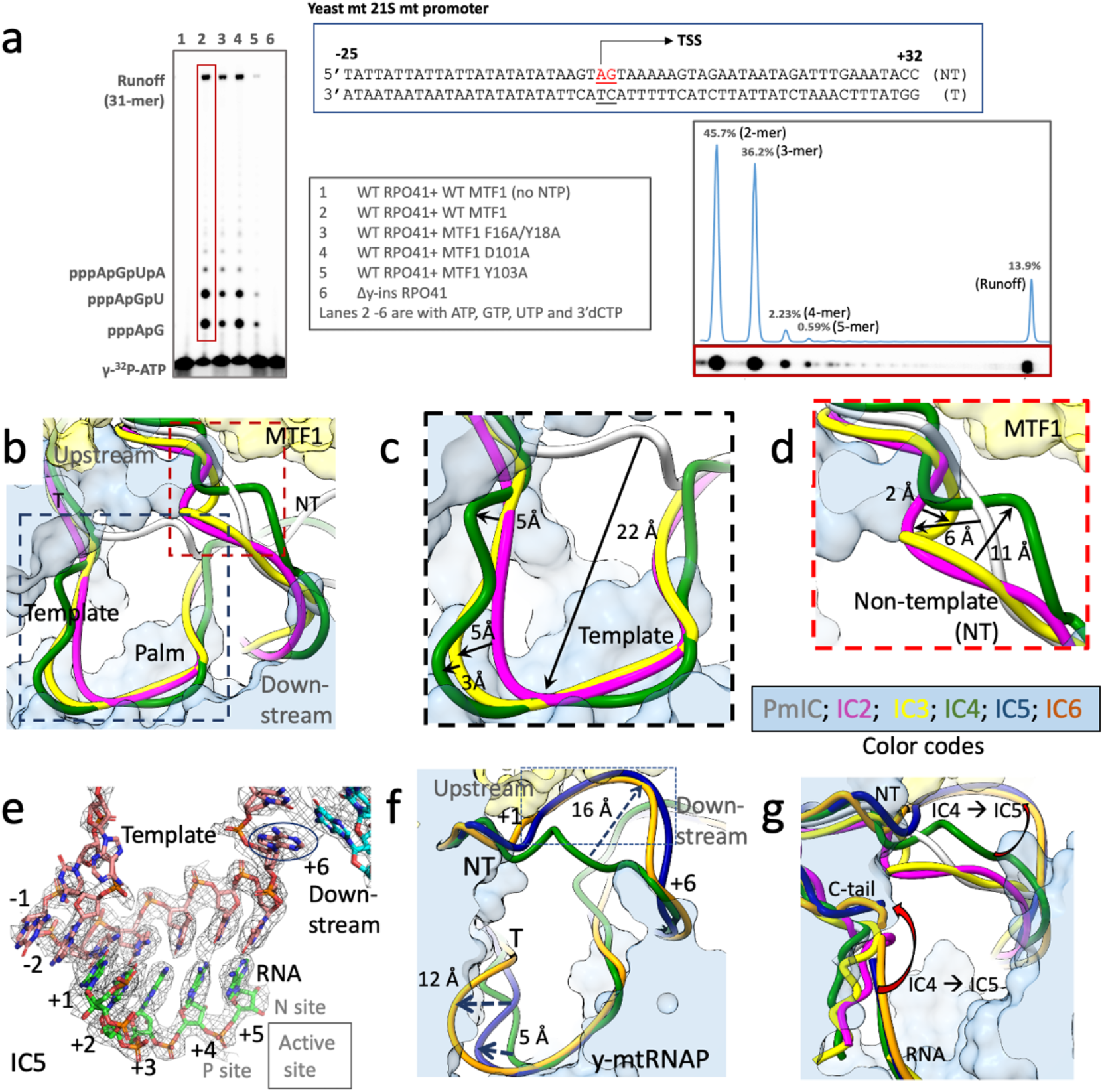
The template, non-template, and MTF1 C-tail in the transcription bubble assume different conformations in different IC states. **a**. The gel image and quantitation of a transcription runoff assay on a consensus y-mtRNAP promoter show that the RNA synthesis is processive after highly abortive 2 -and 3-mer synthesis. **b**. The conformations of template and non-template (NT) strands in the transcription bubbles of PmIC (gray), IC2 (magenta), IC3 (yellow), and IC4 (green). The NT chains are clustered at the top and templates are at the bottom. **c**. Zoomed view showing a stepwise scrunching at the template -1 position as the structure moves from PmIC ⟶ IC2 ⟶ IC3 ⟶ IC4. **d**. Zoomed view showing the scrunching of the NT strand into loops in IC2 and IC3 and then relaxed in IC4. **e**. Experimental density at 3α showing the 5-bp RNA:DNA in IC5 and the unpaired 6^th^ template base (in blue circle) is poised to translocate into the active site cleft for engaging the next nucleotide during IC6 formation. **f**. The transcription bubble conformations in IC4 (green), IC5 (blue), and IC6 (orange); the NT strand has a significant conformational switching in IC4 ⟶ IC5 transition, and the template strand scrunches the most in IC6. **g**. The positioning and conformation of MTF-1 C-tail (bottom) with respect to the NT strand (top) in different IC states, from PmIC to IC6; the red arrows show the conformational switching of both the C-tail and NT strand at IC4 ⟶ IC5 transition.

What triggers the significant dissociation of 2 -and 3-mer RNAs compared to 4-mer or longer RNAs during transcription initiation? Studies of bacterial RNAPs show template scrunching and steric clash with structural elements in the polymerase cleft create stressed intermediates that release RNA products and return to relaxed states, aborting RNA synthesis^28-30^. In y-mtRNAP ICs, we observed distinct positions, conformations, and interactions of the template and NT strands as the transcription bubble grows with each nucleotide addition. The template and NT strands reside in the polymerase cleft and share space with the MTF1 C-tail. Starting from IC2, the template DNA is organized into a U-shaped structure in the polymerase cleft (Fig. 3b). This U-shaped template structure is conserved in T7 RNAP IC^21^, suggesting a standard template track in single-subunit RNAP ICs. As the RNA:DNA hybrid grows in size from IC2 ⟶ IC3 ⟶ IC4, the single-strand region of the template gets scrunched and bulges at the -1 position (Fig. 3c). A scrunched state is known to hold energy and has a tendency to unscrunch to release the stored energy, which destabilizes the RNA:DNA hybrid and releases short RNAs. Thus, template unscrunching could be responsible for 2 -and 3-mer RNA dissociation. However, the template is scrunched the most in IC4 among the three states, yet IC4 is significantly less abortive (Fig. 3a), suggesting that template scrunching is not the primary driving force for abortive synthesis of 2 -and 3-mer RNAs. Our structures show the NT strand is uniquely scrunched in IC2 and IC3 states into tight loops (Fig. 2d, 3d). The scrunched NT loops in IC2 and IC3 are stabilized by interacting with y-mtRNAP intercalating hairpin (641-642) and thumb (780-787), and MTF1 C-tail residues (334-336). We argue that the NT loop unwinding releases energy, like the unwinding of a watch spring, and reverts IC2 and IC3 to the PmIC state by dissociating 2 -and 3-mer RNAs, respectively (Fig. 2c). Interestingly, deletion of the MTF1 C-tail reduces the relative amount of 3-mer abortive, presumably by decreasing DNA scrunching and providing more space for the growing bubble^31^. Overall, the y-mtRNAP IC structures are the first to provide structural evidence involving non-template strand scrunching in the abortive synthesis of short RNAs.

We expected the NT loop to grow bigger with RNA synthesis. However, the IC4 structure shows a relaxed NT strand, which is less ordered with relatively poor density (Supplementary Fig. 1) and a high average B-factor as the strand lacks significant interactions with y-mtRNAP or MTF1. The flexible NT strand holds less stressed energy. Overall, the structural features of IC4 – (i) the half-translocated state of RNA:DNA, (ii) the partially melted +5 base pair in the downstream DNA (Fig. 2g), and (iii) less stressed NT strand support RNA extension over the release of 4-mer RNA.

### Non-template strand forms an ordered base-stacked structure in IC5 and IC6

The IC5 is captured with a 5-bp RNA:DNA hybrid in the pre-translocated state with the RNA 3’-end at the N site. Unexpectedly, the downstream +6 base pair has already melted in IC5. This facilitates IC5 ⟶ IC6 transition (Fig. 3e); upon the availability of the net nucleotide, the unpaired +6 template nucleotide readily flips to engage the next NTP at the N site and extends the RNA:DNA duplex to 6-bp. The growing RNA:DNA and template scrunching in IC5 and IC6 drives several unexpected conformational changes in the MTF1 C-tail and NT strand. These changes stabilize the intermediates and efficiently drive RNA synthesis from 4-mer ⟶ 5-mer ⟶ 6-mer with negligible abortive RNA products compared to 2 -and 3-mer RNAs (Fig. 3a). The growing RNA:DNA continues to scrunch the template strand (Fig. 3f), impacting the MTF1 C-tail (Fig. 3g). In IC2 – IC4, the C-tail is positioned between the RNA:DNA duplex and the thumb subdomain. As the RNA:DNA grows to 5-bp in length, potential steric conflict with the bulged template flips the C-tail towards the center of the transcription bubble. The C-tail flipping causes an unexpected ∼16 Å repositioning of the NT strand from the polymerase cleft center in IC4 to the N-terminal domain of MTF1 in IC5 (Fig. 3f, g). An earlier study has shown that C-tail deletion delays the IC ⟶ EC transition by two nucleotide addition steps^31^. About 10 residues of the MTF1 C-tail occupy the central part of the transcription cleft, and in IC6, this cleft is almost fully occupied by the RNA:DNA duplex, expanded NT, MTF1 C-tail, and the bulged template (Fig. 1e). Hence, deleting the C-tail will create space to accommodate two additional nucleotides in the transcription bubble, delaying the transition to EC.

The repositioned NT strand in IC5 gets organized as a “spiral staircase” like structure even in the absence of a complementary strand (Fig. 4A). The unexpected melting of the +6 base-pair downstream in IC5 is driven by the participation of the NT +6 base in the stacked staircase structure. The NT +1 to +6 nucleotides in the staircase-like structure are highly ordered and well-resolved in the cryo-EM density map, and the stacking remains unchanged during the IC5 ⟶ IC6 transition. The aromatic-ring stacking of the NT bases extends upstream from the +1 position to MTF1 Y103 and the NT (−2)G base.

**Fig. 4.**
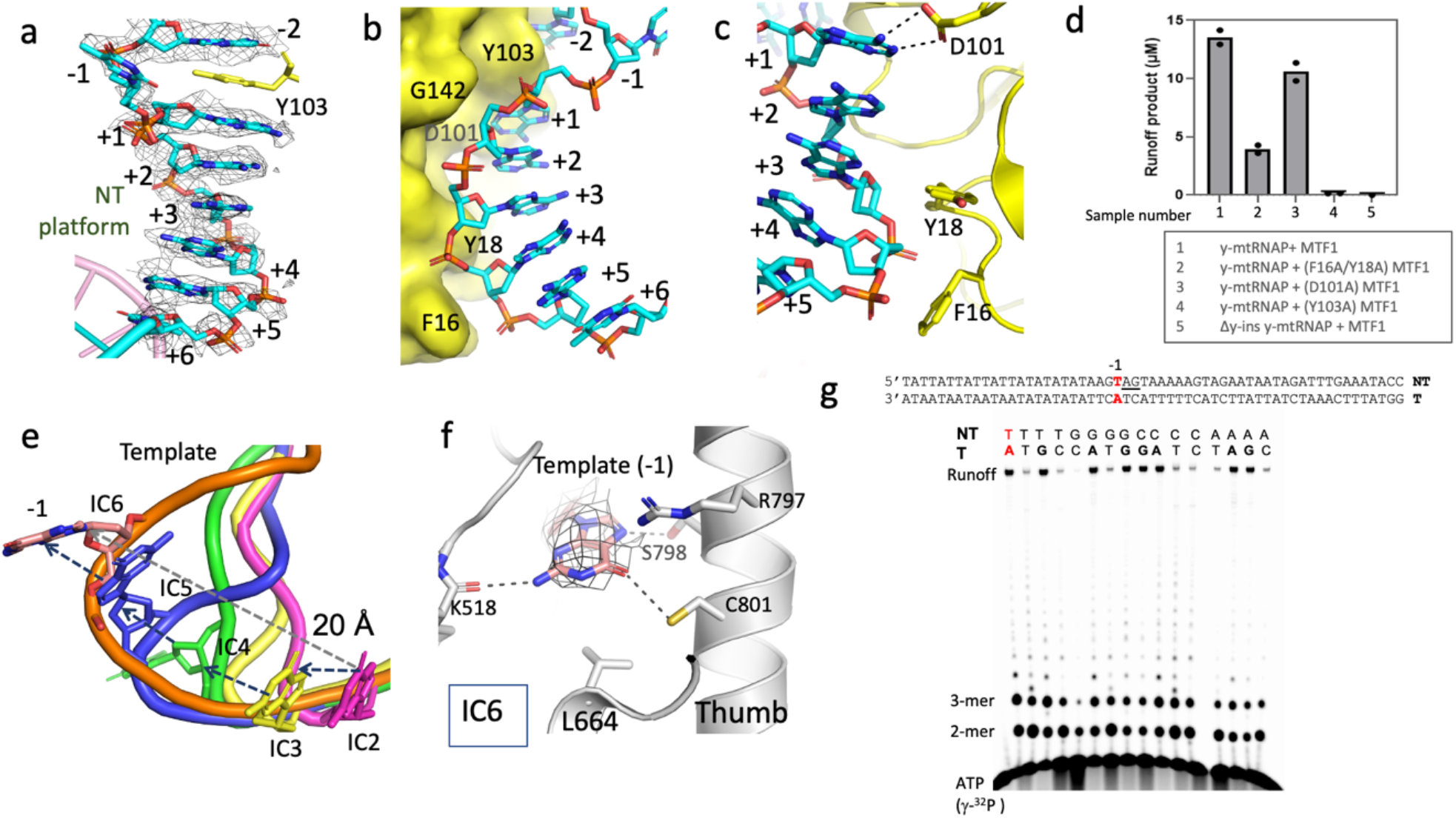
Unexpected structural rearrangements to reach the IC5 and IC6 states, and role of a purine base at template -1 position. **a.** The NT +6 to +1 bases are stacked to form a spiral staircase-like structure in IC5 and the stacking extends upstream to the position -2 NT base involving the aromatic side chain of Y103; the experimental density is displayed at 3σ. **b.** The sugar-phosphate backbone of the stacked NT bases in the spiral staircase is walled against the N-terminal domain of MTF1, shown as yellow molecular surface. **c.** Another view (∼180º of b) of the spiral staircase shows the specific interactions of the NT stack with the D101 sidechain and the F16/Y18 groove. D101 forms salt bridge interactions with AMP at +1 position, and the F16/Y18 groove interacts with NT +4 /+5 sugar-phosphate backbone. **d.** The RNA products shown in the gel image (Fig. 3a) are quantified on the graph and compared with runoff efficiencies of the mutants with wild-type initiation complexes. The amount of runoff product is micromolar. **e.** Relative positions of the template -1 nucleotide in IC2 to IC6 show the progression of the template scrunching with each nucleotide addition. The -1 nucleotide moves ∼20 Å in the transition from PmIC to IC6. **f.** The template position -1 guanine base intrudes into a pocket and interacts with the indicated thumb residues. **g.** The gel image shows *in vitro* transcription runoff assays on DNA promoters substituted with different nucleotides at the NT and template -1 position. The RNA bands were quantified to determine the relative amounts of 2-mer, 3-mer, runoff, and total products (Supplemental Fig. 5).

The ordered NT structure is supported by backbone interactions with the MTF1 N-terminal domain (Fig. 4b, c). The +4/+5 sugar-phosphate backbone of the stacked NT staircase interacts with the F16/Y18 groove of MTF1, and F16A/Y18A mutant MTF1 decreases initiation and runoff RNA products (Fig. 4d). The effect is because of the role of the F16/Y18 groove in the early steps of initiation (Fig. 2e, f). The +1-adenine base in the NT staircase makes salt-bridge interactions with MTF1 D101 (Fig. 4c). Biochemically, substituting the +1 adenine with a 2-aminopurine base that lacks the 6-amino group abolishes runoff transcription but not abortive products^32^, and substituting with 2,6-diaminopurine that contains the 6-amino group rescues the activity to WT promoter level^33^, which suggests the conserved NT +1-adenine in y-mt promoters is essential in later stages of transcription initiation. The MTF1 mutation D101A shows ∼30% drop in runoff product, whereas MTF1 mutation Y103A reduces runoff synthesis by 30-fold (Fig. 4d). This drastic effect of Y103A mutation is expected because Y103 is involved early in the promoter melting step at the PmIC state, and continues to stack with the conserved NT (−2)G base in all IC states. In general, the impact of an altered promoter nucleotide or a protein residue is more drastic at an early than at a late stage of initiation. The base stacking destabilization of the NT staircase is also expected to unsettle the IC states. The +1 to +6 NT sequence in the y-mt promoter is rich in adenines that stack better than thymines^34^. Studies have shown that substituting the NT strand adenines between positions +3 to +8 with a string of thymine bases impairs runoff synthesis, whereas adding 6-12 successive thymines beyond the NT +12 position had little effect^35,36^.

The y-ins domain interacts with the downstream DNA the most in IC5 and IC6 compared to other IC states (Fig. 1f). Biochemically, Δy-ins mutant y-mtRNAP is inactive in transcription initiation (Fig. 3a, 4d), suggesting its greater role in supporting downstream duplex for transcription bubble opening. The stacked bases in IC6 make the NT strand rigid, which may help release upstream DNA contacts as the IC state undergoes a large conformational switching to EC in the next few nucleotide addition steps, as discussed below.

### The template -1 base is critical for the transition from IC to EC

The template scrunching impacts the position of the -1 nucleotide to the maximum (Fig. 3f, 4e). From IC2 to IC6, the -1 template backbone moves by ∼20 Å. In IC2 and IC3, the -1 template base stacks with the RNA:DNA duplex (Fig. 2d), which presumably (i) stabilizes the +1 template:NTP base-pair for *de novo* synthesis of 2-mer RNA, and (ii) stabilizes the 2-mer RNA for the next nucleotide addition to make a 3-mer RNA. Bacterial RNAPs also employ -1 template purine base-stacking to stabilize the RNA:DNA hybrid^37^. In IC4, the -1 template base unstacks and becomes less ordered. In IC5, the -2 base stacks with the RNA:DNA and the -1 base stacks with the -2 base (Fig. 3e). In IC6, the -1 template base unstacks and enters a pocket near the thumb subdomain, where it makes purine-specific interactions with residues of the thumb (R797, S798, C801) and NTD (K518 and L664) (Fig. 4f). Modeling different bases in the template -1 pocket suggests loss of interactions for pyrimidines (Supplementary Fig. 4); i.e., IC5 ⟶ IC6 ⟶ IC7 transitions might be impacted when -1 template is a pyrimidine. A previous report showed that mutation of thumb domain residues, including R797 and R800 increases abortive and termination products after 6-mer synthesis^38^.

To understand the role of -1-template base stacking in IC2/IC3 and later in IC6, we substituted the -1 NT and template nucleotides in a duplexed 21S promoter of y-mtRNAP with all four bases (A/G/T/C), creating matched and mismatched -1 base pairs. These promoters were tested in transcription runoff assays (Fig. 4g). Substituting template -1-adenine with guanine has little impact on the runoff RNA yield; the bubble promoters used in our current cryo-EM study have guanine at the template -1 position. In contrast, substituting -1-adenine with thymine or cytosine reduced the runoff RNA yield by 65 to 600-fold (Supplementary Fig. 5). High-resolution gel analysis of the RNA products shows all promoters are competent in 2 -and 3-mer synthesis, but RNA elongation beyond this length is impaired for promoters with a pyrimidine at -1 template position. Promoter with an abasic -1 template shows 2-mer synthesis, but little 3-mer and no runoff synthesis (Supplementary Fig. 5). Quantifying total RNA products indicate that efficient initiation and synthesis beyond 3-mer requires a pyrimidine in the NT and a purine at template -1 position.

### Modeling the transition from IC to EC

The hallmarks of IC to EC transition are (i) dissociation of the upstream DNA from the RNAP, (ii) reannealing of the -4 to -1 bases to collapse the bubble upstream of RNA:DNA, and (iii) formation of an RNA-exit channel. Previous single-molecule FRET studies of y-mtRNAP showed abrupt unbending/unscrunching of the DNA at 8-nt RNA synthesis, corresponding to upstream promoter release in the transition from IC to EC^4^. This differs from T7 RNAP, where the transition happens through a series of conformational changes between 9-12 nucleotide synthesis^27,39^. Thus in y-mtRNAP, significant structural changes are expected in a span of successive two nucleotides (7^th^ and 8^th^) addition steps. Based on the shifts of the template between the IC2 to IC6 structures (Fig. 4e), we predict the longer RNA:DNA hybrids and highly bulged templates in IC7 and IC8 clash with the MTF1 C-tail to disrupt upstream promoter contacts. The NT staircase is expected to expand and destabilize MTF1 interactions with the NT (−4) AAG (−2) bases to further assist in upstream promoter release. We attempted to obtain an IC7 structure by adding GTP to the reaction following IC6 (Fig. 1a). However, we captured the IC6 state only and not the IC7, suggesting that IC7 is an energetically unstable transient intermediate in the process of switching to EC. We did not observe a backtracked complex predicted by previous single-molecule FRET studies^4^. Unlike DNA unbending, the initial bubble collapse is not abrupt in y-mtRNAP and instead occurs between 8-to 10-nucleotide synthesis^31^. Thus in y-mtRNAP, the IC6 may be the final stable state before the complex undergoes significant structural rearrangement to release the upstream DNA and reanneal the -4 to -1 nucleotides to form the transcription elongation bubble (Movie 1).

The non-reversible conformational changes in y-mtRNAP during the IC ⟶ EC transition are expected to be analogous to those observed for T7 RNAP and h-mtRNAP^23,39^. The T7 RNAP IC8 superimposes well on y-mtRNAP IC6 with nearly aligned nucleic acid tracks (Supplementary Fig. 6). The downstream DNA and RNA:DNA in y-mtRNAP IC6 and h-mtRNAP EC^23^ align very well based on the mtRNAP superposition. Thus, the RNA:DNA and the downstream DNA tracks remain relatively unaffected in the transition from IC to EC (Fig. 5a). However, the upstream DNA is expected to undergo a large swivel motion of ∼110º to assume a relaxed EC state. The scrunched -1 template is the pivot point where the released upstream DNA would rotate to switch its track from IC to EC (Fig. 5b), while the RNA:DNA duplex and the stacked NT platform would remain relatively unchanged. This switching of upstream DNA from IC to EC predicts a major clash with the C-terminal domain of MTF1 supported by the MTF1 hairpin. Interestingly, the MTF1 (or TFB2M) hairpin of y-mtRNAP (or h-mtRNAP) that supports the C-terminal domain of MTF1 in IC structures is engaged in supporting the upstream DNA in EC (Fig. 5c, d).

**Fig. 5.**
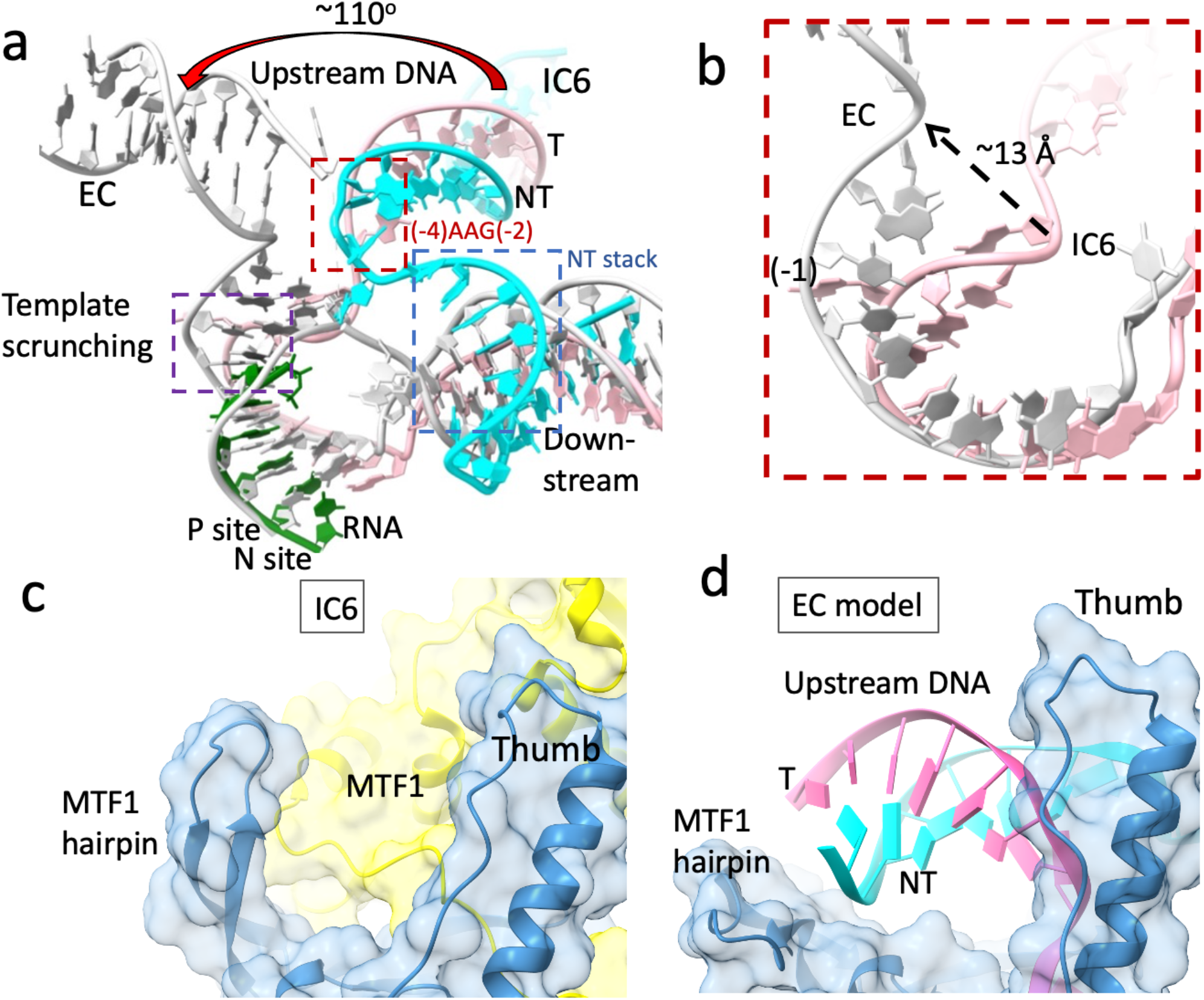
Large structural change associated with the transition from IC6 to EC. **a.** Superposition of y-mtRNAP IC6 and h-mtRNAP elongation complex (PDB: 4BOC; gray) ^23^ by aligning the respective RNAPs shows the common tracks of downstream DNA and RNA:DNA duplex, and the flipping of the upstream DNA as IC6 progresses to EC. The destabilization of NT (−4)AAG(−2) interaction with MTF1 is expected to trigger the flipping of the upstream DNA by ∼110º. The scrunched -1 template position acts as the pivotal point for the upstream DNA flipping. **b.** A zoomed view shows the structural rearrangement at the template -1 region in the IC6 ⟶ EC transition. **c.** In the IC structures, the MTF1 hairpin of y-mtRNAP supports the C-terminal domain of MTF1. **d.** In EC, the MTF1 hairpin switches its role and accommodates the upstream DNA.

## Discussion

In this study, we visualize the entire transcription initiation process from promoter melting to 6-mer RNA synthesis at single nucleotide addition steps (Movies 1 and 2). The structures show that the interactions of y-mtRNAP and MTF1 with the upstream promoter DNA remain intact throughout initiation. The fully resolved transcription bubbles show that the flexible elements of IC, including the NT strand and the MTF1 C-tail, are constantly rearranged in unexpected ways to accommodate the growing RNA:DNA and help stabilize the scrunched IC states. The single-stranded region of the template scrunches continuously, driving changes in the NT and MTF1 C-tail. Template scrunching appears critical for driving the transition from IC to EC. In y-mtRNAP, abortive synthesis of 2 -and 3-mer RNAs is significantly high. This correlates with NT strand scrunching that occurs uniquely in IC2 and IC3. After IC3, RNA synthesis becomes less abortive. The IC5 structure reveals an unexpected strategy to stabilize the expanding NT strand. Instead of scrunching, the +1 to +6 bases of the NT strand stack into a spiral staircase to adapt an ordered low-energy state in IC5. This conformational switching of the NT strand is driven by C-tail repositioning during the IC4 ⟶ IC5 transition. The spiral staircase DNA structure stabilizes the late IC states and drives 4-nt ⟶ 5-nt ⟶ 6-nt elongation.

The NT strand stacking could be a general strategy for creating low-energy states to elongate the RNA at later stages of initiation and subsequently assist in the transition into elongation. Few structures of ICs in bacterial systems have captured *de novo* RNA synthesis on a transcription bubble^37,40^. The NT strands in those structures are either at low resolution or not fully resolved. In bacterial initiation complexes, the purine-rich discriminator element of a promoter, located immediately downstream of the -10 element, has stacked bases like the +1 to +6 NT bases of y-mt promoters. However, the discriminator NT stack is formed during promoter melting and helps with TSS selection^41^. In y-mtRNAP, +1 to +6 NT stacking is formed after 4-mer RNA synthesis, stabilizes the system at the later stage of initiation, and likely guides IC ⟶ EC transition.

The initiation mechanism of h-mtRNAP is expected to be analogous to y-mtRNAP. Both y-mtRNAP and h-mtRNAP form a small initiation bubble and rely on MTF1 and TFB2M, respectively, for promoter melting^31,42,43^. TFB2M is structurally and functionally homologous to MTF1, and biochemical studies have shown that the C-tail of TFB2M has a similar role as MTF1 C-tail in template alignment and initiating NTP binding. Deleting the TFB2M C-tail has a more drastic impact on the initiation and abolishes runoff synthesis. Thereby, many of the processes observed in y-mtRNAP ICs would hold true for h-mtRNAP ICs, but there are differences as well. The h-mtRNAP requires an additional factor TFAM to position and activate the h-mtRNAP:TFB2M at the promoter start site ^44,45^. The y-mtRNAP does not need ABF2, the TFAM homolog, to catalyze promoter-specific initiation, presumably because y-mt promoters contain a consensus sequence unlike the h-mtRNAP promoters that are less conserved. Additionally, y-mtRNAP transcription initiation involves an essential y-ins domain that is absent in h-mtRNAP. As shown here, the y-ins has a critical role early in positioning the promoter DNA and later in stabilizing the downstream DNA (Fig. 1f); this task is apparently delegated to TFAM in h-mtRNAP. With some critical differences, most transcription initiation steps by single-subunit RNAPs are expected to be conserved. In the y-mtRNAP initiation, PmIC is an interesting and unexpectedly stable state. It will be interesting to see if h-mtRNAP also has an analogous PmIC state.

The general principles of transcription initiation mechanism are likely conserved in single-subunit and multi-subunit RNAPs. As proposed for the well-studied bacterial system^7^, our structures show that in single-subunit RNAP (i) the stress-energy is accumulated with unwinding of each downstream base-pair and the bubble expansion, and (ii) an early release of this energy drives the system in an opposite direction leading to release of short abortive products, and (iii) at a later stage the stress-energy pushes the system forward to release the upstream DNA and rotate it about the template -1 nucleotide to enter the elongation phase. The transcription initiation process of the more complex nuclear RNAPs rely on multiple transcription factors for regulation compared to single-subunit and bacterial RNAPs, where the IC states are largely stabilized by structural rearrangements.

## Methods

### Expression and purification of MTF1 WT and mutants

pTrcHisC plasmid encoding His-6 tagged *Saccharomyces Cerevisiae* MTF1 was transformed into *Escherichia coli* BL21 competent cells and expressed and purified as described previously^22,32^. Cells were grown at 37°C in LB media supplemented with 100 µg/mL ampicillin to an OD_600_ of 0.8 before induction with 1 mM iso-propyl β-D-1-thiogalactopyranoside (IPTG). After 16 h at 16°C, cells were harvested and the cell pellet was lysed by sonication in the presence of protease inhibitor and lysozyme in lysis buffer (50 mM sodium phosphate pH 8, 300 mM NaCl, 10% glycerol, 0.1 mM PMSF), and subsequently subjected to polyethyleneimine (10%) and ammonium sulfate (55%) precipitation. The dissolved ammonium sulfate pellet was loaded on a 5 mL DEAE Sepharose and 5 ml Ni-Sepharose cartridge (GE Healthcare Life Sciences) connected in tandem. After sample loading, the DEAE Sepharose cartridge was detached and the Ni-Sepharose cartridge was washed with 50 mL wash buffer (50 mM sodium phosphate buffer pH 8, 300 mM NaCl, 10% glycerol, 1 mM PMSF, 20 mM imidazole). MTF1 protein was eluted with a 70 ml gradient of 20 mM to 500 mM imidazole in wash buffer. The MTF1 peak eluent was collected and loaded on 2 × 1 mL Heparin-Sepharose columns (GE Healthcare Life Sciences) connected in tandem. Subsequently, the columns were washed with 20 ml heparin wash buffer (50 mM sodium phosphate buffer pH 8, 150 mM NaCl, 10% glycerol, 1 mM EDTA, 1 mM DTT, 1 mM PMSF) prior to elution with a 50 mL gradient of 150 mM to 1 M NaCl in heparin wash buffer. The MTF1 protein eluent was collected and concentrated using 10 kDa MW cut-off Amicon ultra-centrifugal filter (Merck Millipore) and stored at -80°C. The MTF1 mutant proteins were purified using the same protocol (Supplementary Fig. 7).

### Expression and purification of ΔN100 y-mtRNAP and y-ins (1237-1321 deleted) y-mtRNAP

ProEXHTB plasmid encoding His-6 tagged *Saccharomyces Cerevisiae* ΔN100 y-mtRNAP was transformed into *Escherichia coli* BL21 RIL Codon Plus competent cells and expressed and purified as described previously<u>^22,32,46^</u>. Cell growth conditions, lysis, PEI and ammonium sulphate precipitation, tandem DEAE and Ni column, and heparin column chromatography steps were identical to MTF1 described above. The ΔN100 y-mtRNAP protein eluent from heparin column was treated with TEV protease at 100:1 (w:w) ΔN100 y-mtRNAP to TEV protease for 16h at 4°C. The cleaved protein was loaded on a 5 mL Ni-Sepharose cartridge and the flow-through was collected and concentrated using 10 kDa MW cut-off Amicon ultra-centrifugal filter (Merck Millipore) and stored at -80°C. The y-ins deleted y-mtRNAP was purified using the same protocol (Supplementary Fig. 7).

### In vitro transcription initiation assay to measure runoff and abortive RNA synthesis

In vitro transcription initiation assays were carried out, as discussed previously ^22^. Transcription reactions on 2 µM pre-melted initiation bubble promoter was carried out with y-mtRNAP (1µM) and MTF1 (2 µM) in reaction buffer (50 mM Tris acetate pH 7.5, 100 mM potassium glutamate, 10 mM magnesium acetate, 1 mM DTT, 0.01% Tween 20) at 25°C using 20 µM of indicated NTPs spiked with α-^32^P-GTP. RNA synthesis was terminated after 15 min using 125 mM EDTA and formamide dye mixture (98% formamide, 0.025% bromophenol blue, 10 mM EDTA), and RNA products were resolved on a 24% polyacrylamide, 4M urea gel. Visualization, runoff RNA product quantification and analysis of the gel was accomplished with ImageQuant software. Transcription reactions on duplex promoters (1 µM) were carried out under similar conditions, except NTP concentrations were 100 µM and reactions were spiked with γ-^32^P ATP.

### Assembly and characterization of y-mtRNAP IC2, IC4, IC5, and IC6

Complex assembly and characterization were carried out as described previously ^19,22^. Two y-mtRNAP PmICs were prepared by incubating ΔN100 y-mtRNAP, MTF1 and promoter DNA in a molar ratio of 1:1.2:1.2 for two hours at 4°C. The PmIC with the 33-bp bubble promoter was used for y-mtRNAP IC2 and IC4 structures, and PmIC with the 36-bp bubble promoter was used for IC5 and IC6 structures. Both PmICs, at a starting concentration of 6 mg/ml in buffer A (50 mM Bis-tris propane pH 7.0, 100 mM NaCl, 5 mM MgCl_2_, 1 mM EDTA, 2 mM DTT), were subjected to size-exclusion chromatography using a Superdex 200 Increase 10/300 GL column connected to an AKTA PURE 25 FPLC system (GE Healthcare Life Sciences) maintained at 6° C. The column was in-line with a mini-DAWN multi-angle light-scattering device (MALS; Wyatt Technology), a differential refractive index (dRI; Optilab) measuring device, and a dynamic light scattering (DLS; DynaPro Nanostart) device. After purification, the PmICs were concentrated to 3 mg/ml, aliquoted, and stored at -80°C prior to further use.

The PmIC on the 33-bp bubble promoter was incubated with GTP at a molar ratio of 1:50 to generate IC2, and with pppGpG, UTP and ATP at a molar ratio of 1:3:25:25 to prepare IC4. The PmIC on the 36-bp bubble promoter was incubated with pppGpG and ATP at a molar ratio of 1:3:50 to generate IC5, and pppGpG, ATP, UTP and GTP at a molar ratio of 1:3:50:20:20 to generate IC6. All the ICs were incubated for two hours at 4°C prior to grid preparation.

The complexes were characterized by stand-alone cuvette mode DLS experiments using DynaPro Nanostar (Wyatt Technology) to confirm similar hydrodynamic radius and polydispersity for all IC samples. For the DLS experiments, 8 µL of IC samples at a concentration of 0.5 mg/ml were loaded into disposable cuvettes (Wyatt Technologies) and placed in the sample chamber maintained at 4°C. DLS experiments consisted out of 30 acquisitions for each sample and data was analyzed using Dynamics software (Version 7.10.0.23; Wyatt Technology).

### Cryo-EM grid preparation and data collection

Vitreous grids of y-mtRNAP ICs were prepared on Quantifoil R 1.2/1.3 holey carbon grids (Cu300 or Au300) using a Leica EM GP (Leica Microsystems). The grids were glow-discharged for 45 s at 25 mA current with the chamber pressure set at 0.3 mBar (PELCO easiGlow; Ted Pella). Glow-discharged grids were mounted in the sample chamber of a Leica EM GP at 8°C and 95% relative humidity. Optimized grids for IC2 and IC5 were obtained by spotting 3 µL of the sample at 0.8 and 0.9 mg/ml (50 mM Bis-Tris propane, pH 7.0; 100 mM NaCl, 5 mM MgCl_2_, 1 mM EDTA, and 2 mM DTT) on Cu300 grids, respectively. Sample was incubated on the grids for 30 seconds prior to back-blotting for 12 seconds using two pieces of Whatman Grade 1 filter paper and the plunge-freezing were done by dipping the grids in liquid ethane at temperature of -172°C. Optimized grids for IC4 and IC6 were obtained by spotting 3 µL of sample at 0.7 or 0.8 mg/ml on Au300 grids. The grids were clipped and mounted on a 200 keV Glacios™ cryo-transmission electron microscope (Thermo Fisher) with autoloader and Falcon 3 direct electron detector as installed in our laboratory. High-resolution data sets for y-mtRNAP IC2 and y-mtRNAP IC4 to IC6 were collected on the Glacios using EPU software version 2.9.0 (ThermoFisher Scientific).

Electron movies were collected in the counting mode at a nominal magnification of 190.000x for IC2 yielding a pixel size of 0.76 Å, and at a nominal magnification of 150.000x for IC4, IC5 and IC6 yielding a pixel size of 0.97 Å. The total exposure time was 25.15 seconds for IC2, and 38.41 seconds for IC4, IC5 and IC6 with a total dose of 40 e/Å^2^ for all datasets. All movies were recorded as gain corrected MRC files. The data-collection parameters for all structures are listed in Supplementary Table 1.

### Cryo-EM Data Processing

For each dataset, individual movie frames were motion-corrected and aligned using MotionCor2 ^47^ as implemented in the Relion 3.1 package^48^ and the contrast transfer function (CTF) parameters were estimated by CTFFIND-4^49^. The particles were automatically picked using the reference-free Laplacian-of-Gaussian routine in Relion 3.1. The picked particles were subjected to series of 2D and 3D classifications. The final 3D classification generated a distinct class, and no additional class such as PmIC was detected in the processing of any of the datasets. The final set of particles for each IC was used to calculate gold-standard auto-refined maps, which were further improved by B polishing and CTF refinement. All data processing steps were carried out using Relion 3.1.

### Model building

Data processing yielded 3.47 Å, 3.44 Å, 3.39 Å, and 3.62 Å density maps for IC2, IC4, IC5, and IC6 respectively, and were used to fit the atomic model for respective structures. Previously published PmIC (PDB: 6YMV) and IC3 (PDB: 6YMW) structures were used as references for building the models for y-mtRNAP IC2 and IC4-IC6 structures. Furthermore, the AlphaFold prediction of y-mtRNAP (UniProt ID: P13433) was used for modelling the insertion region (y-ins, residues 1232 to 1328). In addition, the y-ins region of previously published PmIC and IC3 structures have been corrected based on the AlphaFold model. All model building were done manually using COOT^50^. Model building was coupled with iterative rounds of real-space structure refinement using Phenix 1.19.2-4158^51^. In the IC2 final structure, MTF1 is traced from S2 – S340 and y-mtRNAP is traced from I386 to the end residue S1351 with missing stretches 559 – 588 and 1309 -1320. In the IC4 structure, MTF1 is traced from S2 to G341 (end residue) and y-mtRNAP from A407 to S1351 (with missing stretches 559 – 588 and 1311 – 1319). In the IC5 structure, MTF1 is traced from S2 to Y335 and y-mtRNAP from I386 to S1351 (residues 556 – 588 and 1310-1319 missing). In the IC6 structure, MTF1 is traced form S2 to T337 and y-mtRNAP is traced from I386 till S1351 (residues 554 – 588 and 1311 – 1319 missing). For each structure, the DNA and RNA chains are different, and were modeled independently to the respective experimental density maps. All structure figures were generated using PyMol (https://pymol.org/2/), Chimera^52^ and ChimeraX^53^. Movies were generated using Chimera.

## Data availability

The data that support this study are available from the corresponding author upon reasonable request. Certain materials will require materials transfer agreements (MTAs). The coordinates and cryo-EM density maps for IC2, I4, IC5, and IC6 have been deposited under PDB accession codes 8AP1, 8ATT, 8ATV and 8ATW, and EMD-15556, EMD-15662, EMD-15664, and EMD-15665, respectively.

## Acknowledgments

We thank Abhimanyu K. Singh for helpful discussion and Neesha Rajesh Shewakramani for her help with movie making. The study was supported by Rega Virology and Chemotherapy internal grants to K.D. and National Institutes of Health grant GM118086 to SSP.

## Author contributions

Conceptualization: SSP, KD

Methodology: JS, BDW, UB

Investigation: QG, JS, SSP, KD

Visualization: QG, SSP, KD

Funding acquisition: SSP, KD

Project administration: SSP, KD

Supervision: SSP, KD

Writing – original draft: SSP, KD

Writing – review & editing: All authors

## Additional information

Supplementary material available.

## Supplementary Materials

## Movie legends

**Movie 1. The structural changes of the nucleic acid parts showing the transition from PmIC to IC6 at single nucleotide addition steps and then from IC6 to EC**. Morphing between the partially-melted initiation state (PmIC), initiation states (IC2 – IC6), and the elongation state (EC) simulates the conformational changes in promoter and proteins in the initiation complexes and during the transition to the elongation complex. The y-mtRNAP is in blue, MTF1 in yellow, non-template (NT) DNA in cyan, template DNA in pink, incoming NTPs in blue, incorporated RNA in yellow, template position -1 in green, and the template in red for the end states and the structural elements are in gray for the start states in the movie. The transcription bubble opens fully during the transition from PmIC to IC2; the +1 and +2 bases flip towards the polymerase active site and base-pair with the incoming NTPs to initiate *de novo* RNA synthesis. This promoter melting is accompanied by further bending of the downstream DNA by about 60° with respect to the upstream DNA. The promoter DNA maintains stable upstream DNA interactions with the y-mtRNAP and MTF1 and is spatially confined throughout the initiation steps PmIC ⟶ IC6. In the PmIC ⟶ IC2 transition, the transcription bubble fully opens, and the downstream template bases are brought in to align with two initiating GTP molecules at the active site; this transition is accompanied by the non-template (NT) scrunching and the single-strand part of the template taking a “U” shape. In the IC2 ⟶ IC3 transition: (i) the third nucleotide (an UTPαS) binds at the N site and is poised for catalytic incorporation, (ii) the template starts to bulge as the RNA:DNA duplex pushes the single strand template region of -4 to -1 nucleotides, and (iii) the NT loop scrunches further. In the IC3 ⟶ IC4 transition: (i) the 4^th^ nucleotide is incorporated to the RNA strand, (ii) the template bulging expands, however, the stacking of -1 template base (in green) with the RNA:DNA is dissociated, and (iii) the scrunched NT strand is liberated. The single strand parts of the template and non-template in the transcription bubble are significantly less ordered and lack interactions with protein residues. In the IC4 ⟶ IC5 transition: (i) the 5-mer RNA is formed, (ii) the template -2 base is stacked with RNA:DNA on one side and with -1 base on the other side, (iii) surprisingly the NT strand switches its position and conformation to stack +1 to +6 nucleotide bases in an orderly fashion as a “staircase-like” structure, (iv) the +6 base pair is melted from the downstream duplex and the +6 NT base is engaged in the stable NT base-stacked structure, whereas, the +6 template base is ready to enter the polymerase cleft for binding of the next NTP. In the IC5 ⟶ IC6 transition, (i) the template is translocated by one nucleotide while the NT strand has changed a little, (ii) RNA grows to a 6-mer, and (iii) the template -1 adenine base has moved significantly to occupy a pocket adjacent to the thumb subdomain. We obtained IC6 as the last stable state before the complex enters EC; our attempt to trap IC7 resulted in IC6 only. The large conformational changes in the transition from IC6 ⟶ EC are visualized by morphing a y-mtRNAP:RNA:DNA EC structure, modeled based on h-mtRNAP:RNA:DNA EC structure (PDB ID. 4BOC), with the y-mtRNAP IC6 structure. The RNA:DNA and downstream DNA duplexes in both structures superimpose, whereas the upstream DNA is released from its interactions with MTF1 and y-mtRNAP and switches conformation to enter the elongation phase. In the process, MTF1 is released, and the -4 to -1 bases of the template and NT reanneal. The above structural transitions are shown as a single uninterpreted clip in the end.

**Movie 2. The structural changes of y-mtRNAP and MTF1 through the transition from PmIC to IC6 at single nucleotide addition steps and then from IC6 to EC**. The PmIC ⟶ IC2 transition is associated with the sliding of MTF1 over y-mtRNAP to close the polymerase and downstream clefts for holding the downstream DNA that helps the bubble fully open and engage the incoming NTPs for *de novo* initiation. The thumb subdomain also moves with MTF1 maintaining the protein:protein interactions. The clefts gradually open with each nucleotide addition to accommodate the growing transcription bubble, and in IC4 the relative positioning of MTF1 and y-mtRNAP almost returns to that in PmIC. In the IC5 and IC6 states, the MTF1 moves in a lateral direction with respect to y-mtRNAP, which is ∼90º from its closing/opening movements in the PmIC to IC4 transitions (SI Fig. S3). The direction of the movement in IC5 and IC6 appears to be along a path that helps the release of MTF1 between the IC6 ⟶ EC transition over the next few nucleotide additions. The IC6 ⟶ EC transition is expected to associate with large conformational changes of the upstream DNA, release of MTF1, and switching in the role of the MTF1-hairpin that holds the C-terminal domain of MTF1 in IC states to accommodate the upstream DNA in EC. The C-tail in the polymerase cleft plays an important role and undergoes large positional and conformational changes to stabilize the transcription bubble at different IC states.

**Supplementary Table 1.** Single Particle Cryo-EM Data and Structure Analysis Statistics.

|  |  |  |  |  |
| --- | --- | --- | --- | --- |
| **Structure** | IC2<br>(y-mtRNAP; MTF1; dsDNA;<br>2GTP) | IC4<br>(y-mtRNAP; MTF1; dsDNA;<br>pppGpGpUpA) | IC5<br>(y-mtRNAP; MTF1; dsDNA;<br>pppGpGpApApA) | IC6<br>(y-mtRNAP; MTF1; dsDNA;<br>pppGpGpApApApU) |
| PDB ID/EMBD ID | 8AP1/EMD-15556 | 8ATT/EMD-15662 | 8ATV/EMD-15664 | 8ATW/EMD-15665 |
| **Data Collection** |  |  |  |  |
| Data collection date | 18-12-2020 | 15-02-2022 | 24-02-2021 | 18-02-2022 |
| Grid type | Quantifoil R 1.2/1.3 Cu300 | Quantifoil R1.2/1.3 Au300 | Quantifoil R 1.2/1.3 Cu300 | Quantifoil R1.2/1.3 Au300 |
| Number of grids | 1 | 1 | 1 | 1 |
| Microscope/detector | Glacios™ Cryo-TEM /<br>Falcon 3 | Glacios™ Cryo-TEM /<br>Falcon 3 | Glacios™ Cryo-TEM /<br>Falcon 3 | Glacios™ Cryo-TEM /<br>Falcon 3 |
| Voltage (kV) | 200 | 200 | 200 | 200 |
| Magnification | 190,000x | 150,000x | 150,000x | 150,000x |
| Recording mode | Counting | Counting | Counting | Counting |
| Dose (e/Å<sup>2</sup>/frame) | 1.58 | 1.04 | 1.04 | 1.04 |
| Total dose (e/Å<sup>2</sup>) | 40 | 40 | 40 | 40 |
| Number of frames/movies | 40 | 40 | 40 | 40 |
| Total exposure time (s) | 25.15 | 38.41 | 38.41 | 38.41 |
| Pixel size (Å) | 0.76 | 0.97 | 0.97 | 0.97 |
| Defocus range (Å) | -8,000 to -22,000 | -8,000 to -18,000 | -8,000 to -22,000 | -8,000 to -18,000 |
| **Data processing** |  |  |  |  |
| Number of micrographs used | 2,045 | 997 | 602 | 1,627 |
| Number of particles picked | 1,149,870 | 671,969 | 549,988 | 940,973 |
| Number of particles after 2D/3D clean | 309,765 | 296,664 | 192,599 | 275,877 |
| Particles used for final map | 137,631 | 138,730 | 91,298 | 144,092 |
| Map resolution (FSC 0.143; Å) | 3.47 | 3.44 | 3.39 | 3.62 |
| Map sharpening B factor (Å<sup>2</sup>) | -144 | -80 | -140 | -125 |
| **Model fitting** |  |  |  |  |
| Experimental map/model correlation | 0.82 | 0.72 | 0.77 | 0.79 |
| Total number of atoms | 11,229 | 11,153 | 11,161 | 11,283 |
| **Average B factor (Å<sup>2</sup>)** |  |  |  |  |
| Protein atoms | 22.34 | 88.32 | 28.60 | 75.27 |
| Nucleid acid and NTP | 51.85 | 80.43 | 79.50 | 95.37 |
| Clash score | 5.85 | 9.84 | 6.95 | 8.42 |
| Ramachandran plot; favored/outlier (%) | 96.07/0.00 | 97.58/0.00 | 96.64/0.00 | 97.20/0.00 |
| Rotamer outlier (%) | 0.27 | 0.18 | 0.00 | 0.00 |
| RMSD bond length (Å)/bond angle (°) | 0.005/0.762 | 0.006/1.080 | 0.005/1.008 | 0.006/1.023 |

## Supplementary Figures

**Supplementary Fig. 1.**
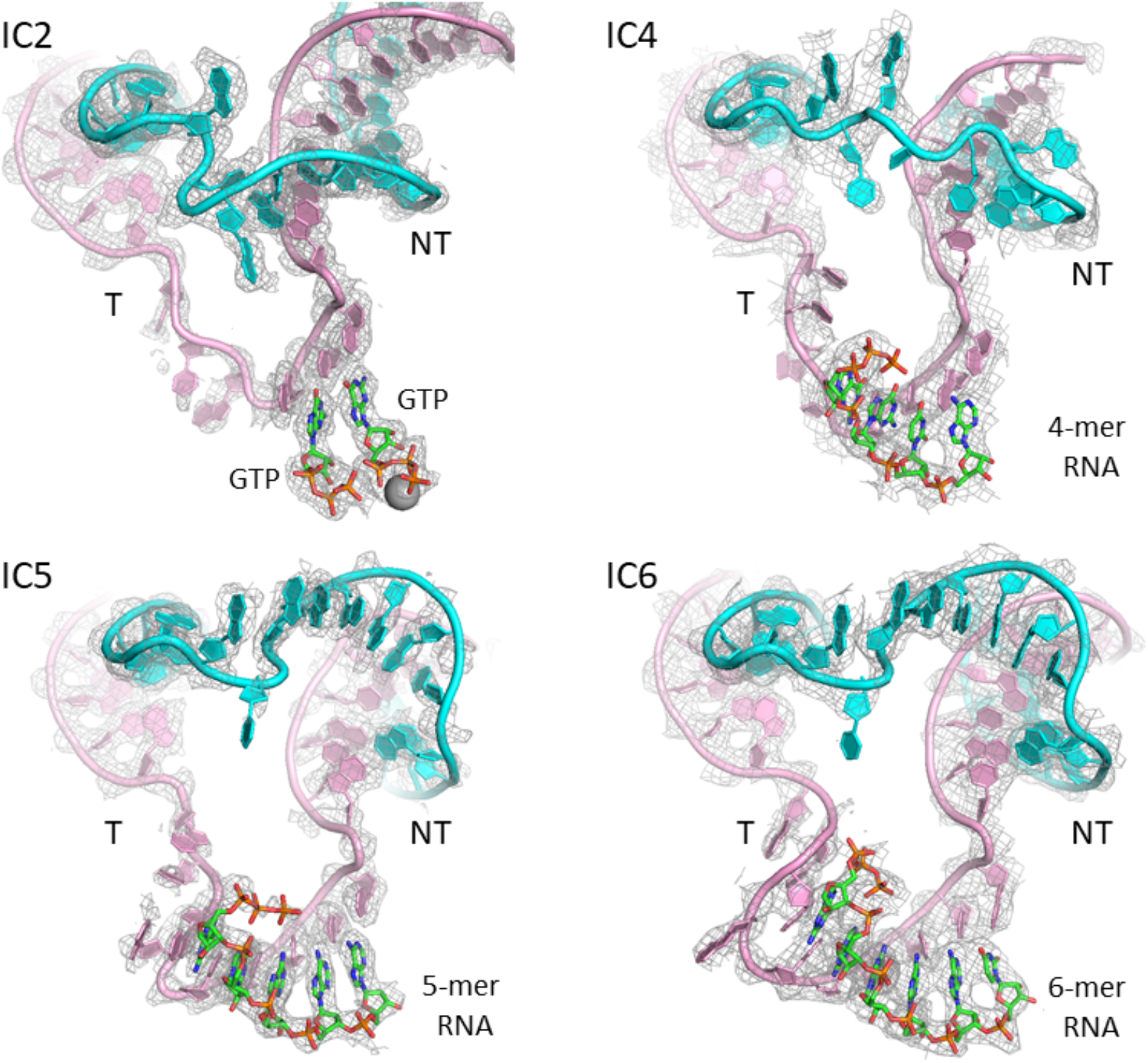
Density maps of the transcription bubbles in IC2, IC4, IC5 and IC6. The density maps clearly define the position and conformation of complete transcription bubbles in different IC structures; with the non-template (NT) in cyan, template (T) in pink, and GTP and RNA in green C-atom representation. The contour levels of the density maps are 1.75, 1.3, 2, and 2σ for IC2, IC4, IC5 and IC6, respectively. The shown density map for IC4 is the unsharpened map as the B-sharpened map is noise. Remaining three maps are B-sharpened.

**Supplementary Fig. 2.**
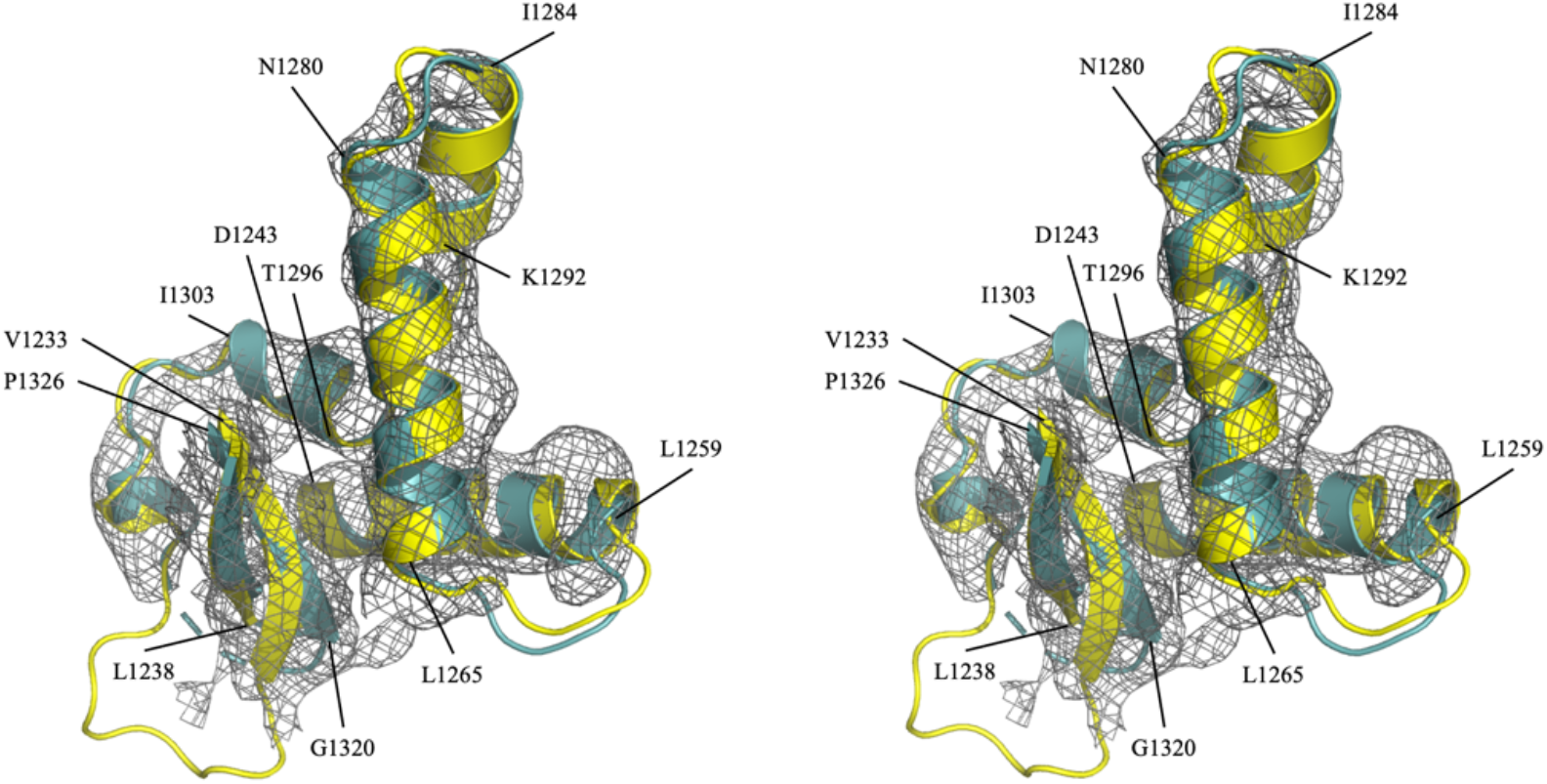
Stereo view of y-ins region. Wall-eyed stereo view showing the starting y-ins AlphaFold model (UniProt ID: P13433 -yellow) that was fitted to the density. The final model (turquoise) after density fit also aligns well with the AlphaFold model; the density segment for the y-ins domain in the IC6 structure is contoured at 2.5σ.

**Supplementary Fig. 3.**
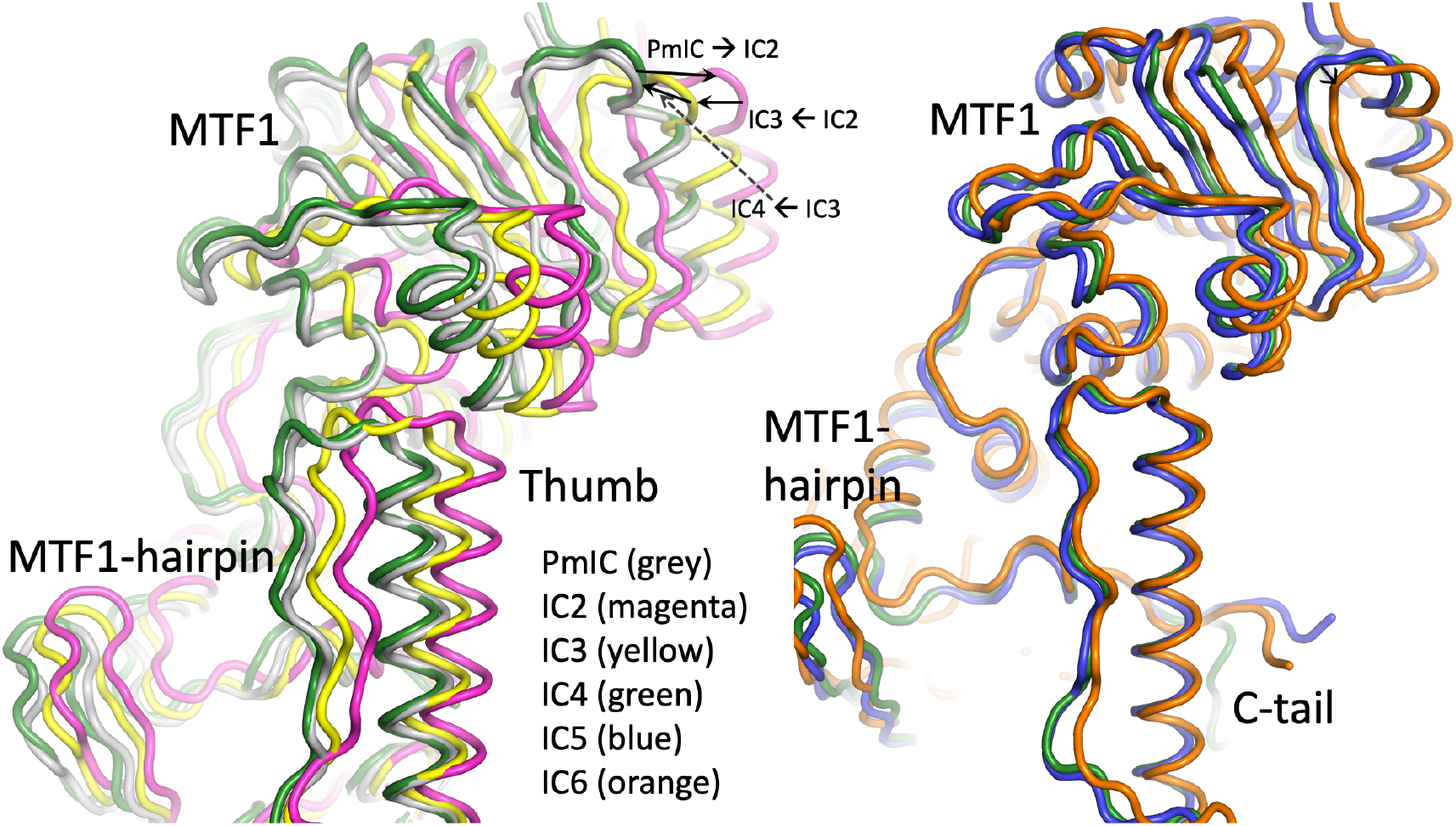
Superposition of MTF1 in different IC states. Positioning of MTF1 and y-mtRNAP (thumb and MTF1-hairpin) in different IC states; from PmIC to IC4 (left) and IC4 to IC6 (right). The structures were aligned based on y-mtRNAP superposition.

**Supplementary Fig. 4.**
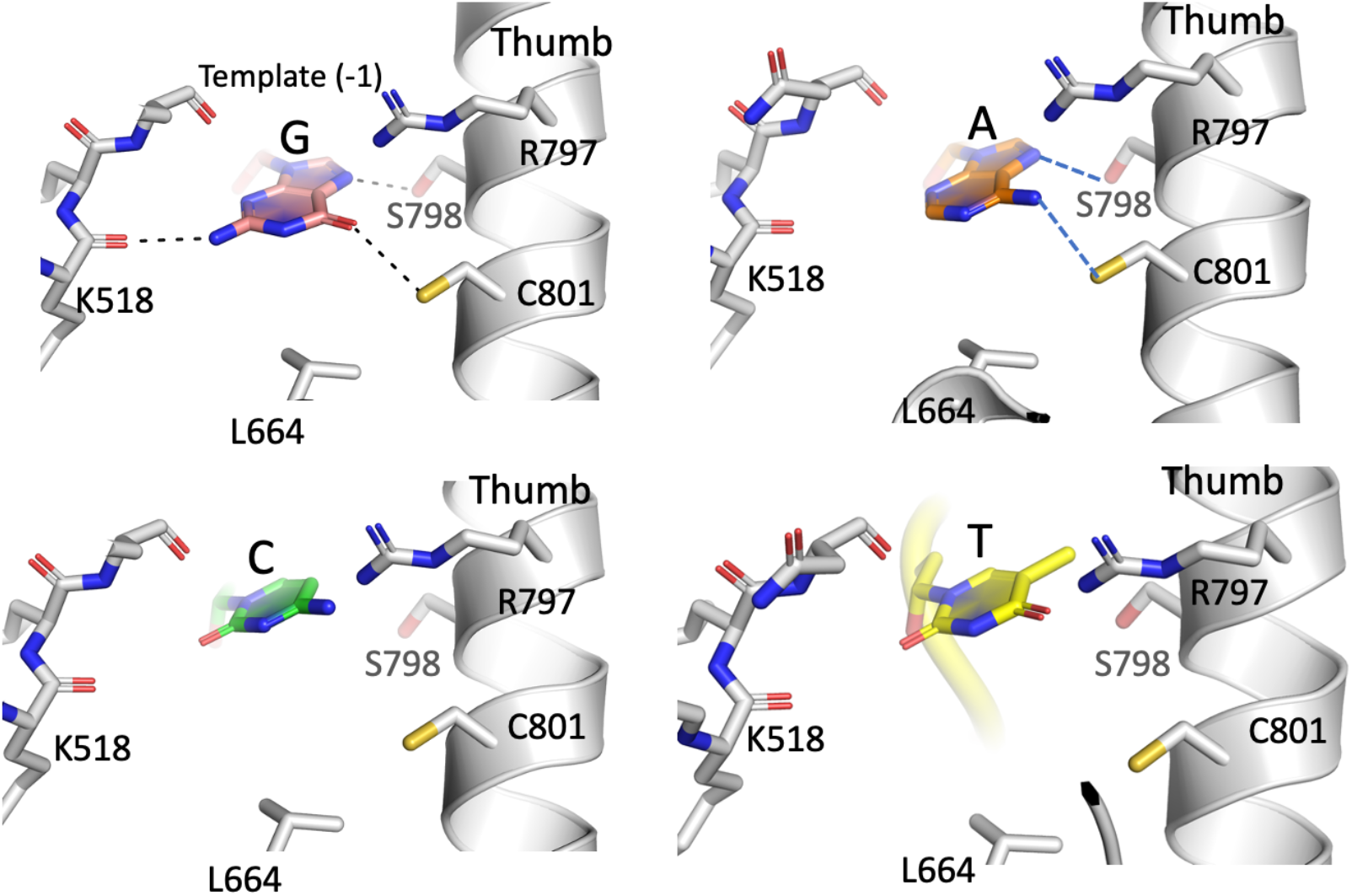
Modeling of A/C/T bases into the template -1 pocket in IC6 structure. Potential interactions of template -1 nucleotide in IC6 structure when a guanine (G) is substituted with adenine (A), cytosine (C), and thymine (T). The base interactions are significantly reduced by a purine to pyrimidine substitution.

**Supplementary Fig. 5.**
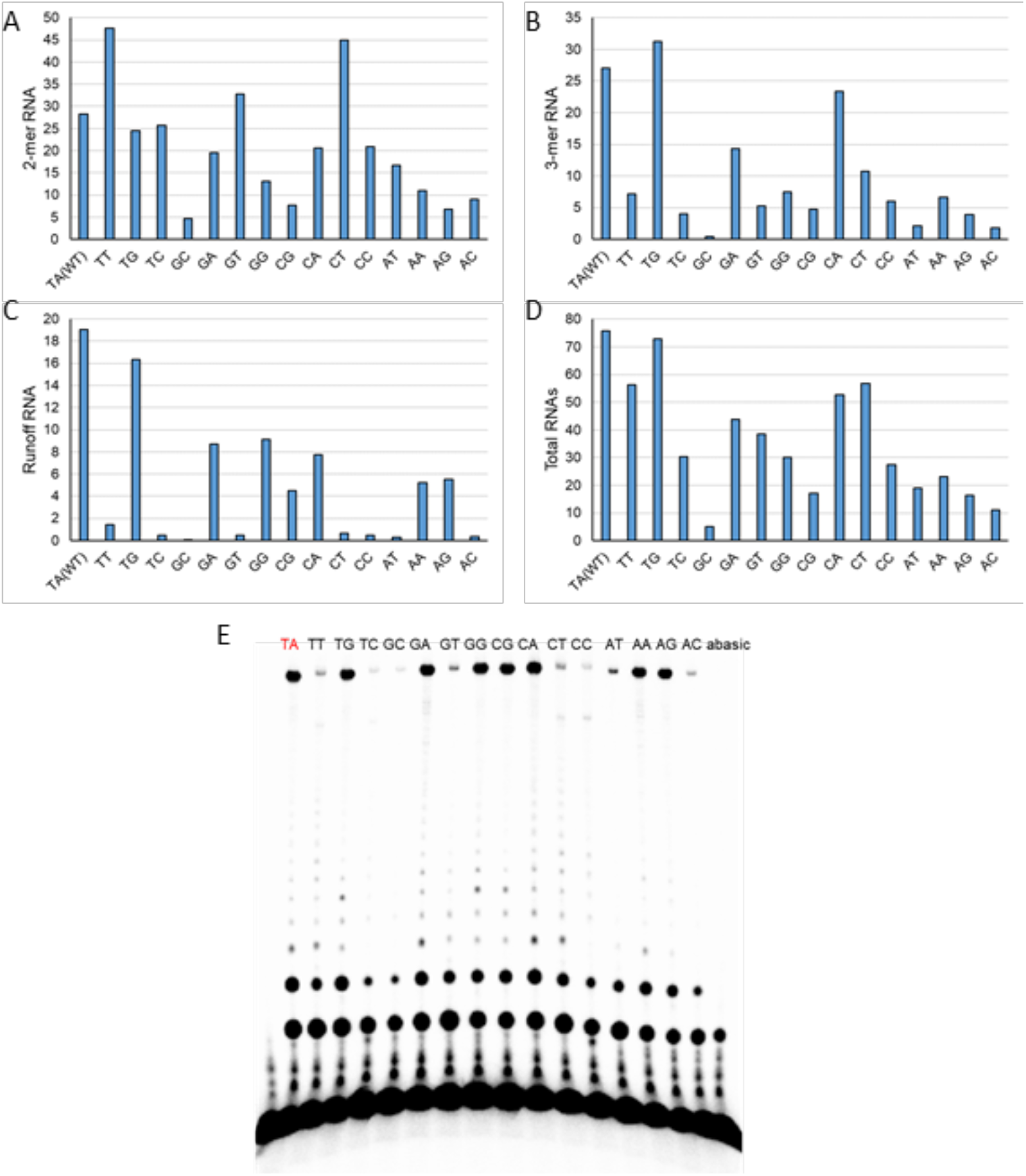
In vitro transcription on -1 position modified promoters. The quantitively analyses of the gel in Fig. 4G show the effect of -1 position changes on 2-mer (A), 3-mer (B), runoff (C), and total RNA synthesis (D). The gel image from a repeated experiment demonstrating reproducibility of the data. An abasic -1 template position was included in this repeat (E).

**Supplementary Fig. 6.**
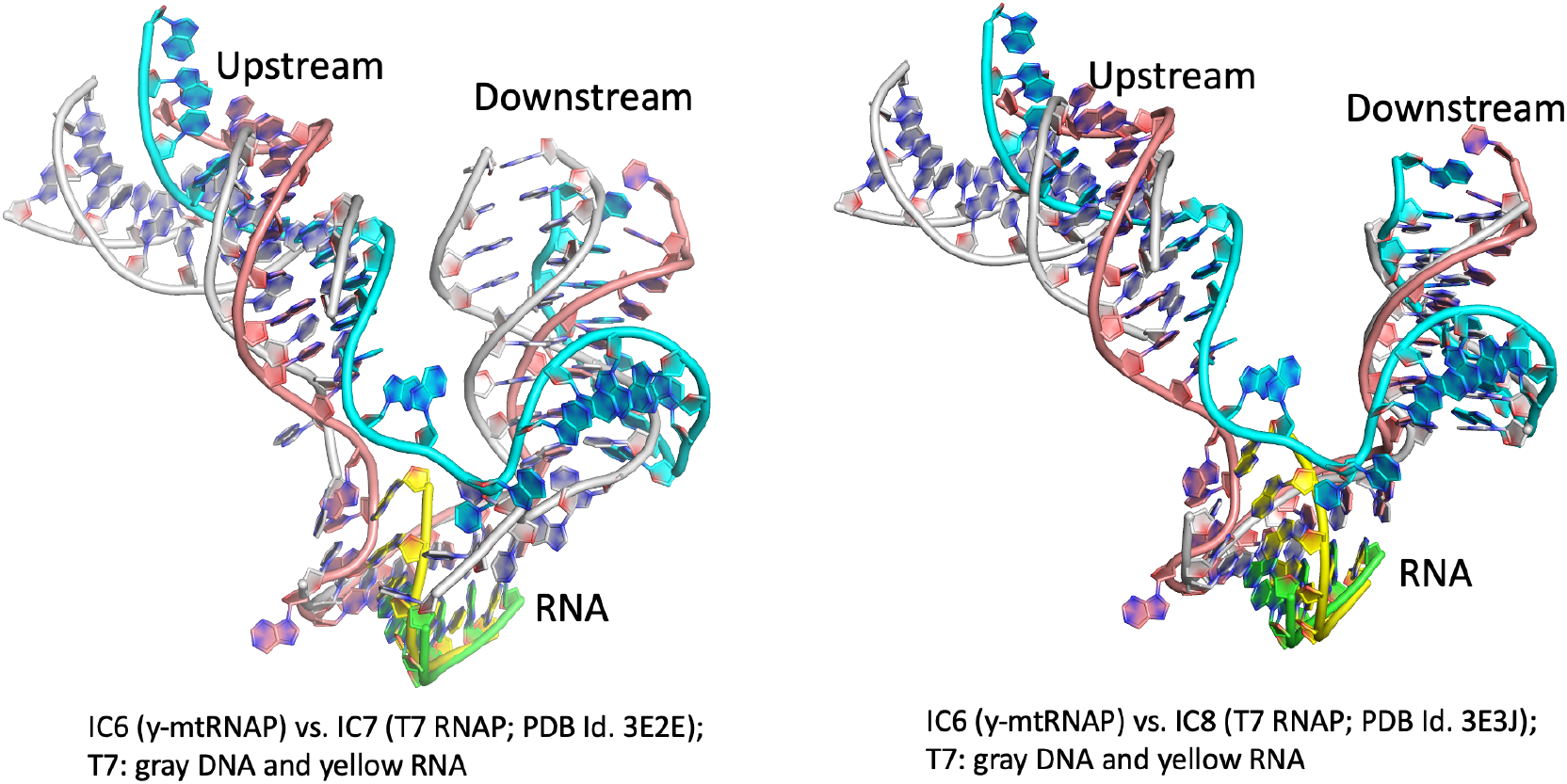
Superposition of IC6 on T7 RNAP IC7 and IC8. Relative positions of upstream and downstream DNA, and RNA:DNA duplex in y-mtRNAP IC6 with T7 RNAP IC7 (left) and IC8 (right). The template, non-template, and RNA in IC6 are colored pink, cyan, and green, respectively. The DNA and RNA in T7 IC7 and IC8 are colored gray and yellow, respectively. The alignments were based on RNAP Cα superposition.

**Supplementary Fig. 7.**
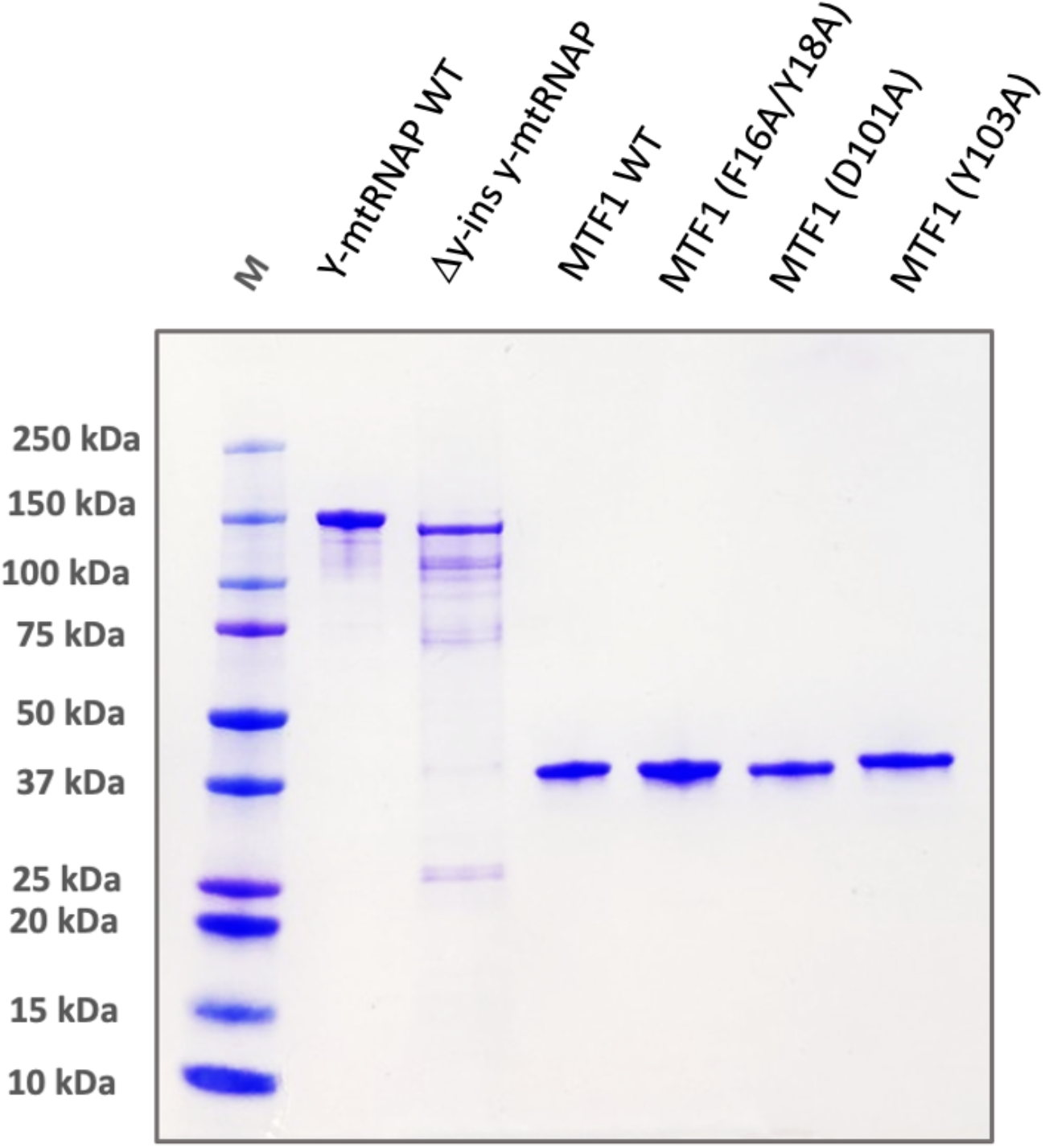
SDS-PAGE analysis of different y-mtRNAP and MTF1 proteins. Purified wild-type and mutant proteins (y-mtRNAP and MTF1) used in transcription runoff assays.

